# eGARD: Extracting associations between genomic anomalies and drug responses from text

**DOI:** 10.1101/148833

**Authors:** A. S. M. Ashique Mahmood, Shruti Rao, Peter McGarvey, Cathy Wu, Subha Madhavan, K. Vijay-Shanker

## Abstract

Tumor molecular profiling plays an integral role in identifying genomic anomalies which may help in personalizing cancer treatments, improving patient outcomes and minimizing risks associated with different therapies. However, critical information regarding the evidence of clinical utility of such anomalies is largely buried in biomedical literature. It is becoming prohibitive for biocurators, clinical researchers and oncologists to keep up with the rapidly growing volume and breadth of information, especially those that describe therapeutic implications of biomarkers and therefore relevant for treatment selection. In an effort to improve and speed up the process of manually reviewing and extracting relevant information from literature, we have developed a natural language processing (NLP)-based text mining (TM) system called eGARD (extracting Genomic Anomalies association with Response to Drugs). This system relies on the syntactic nature of sentences coupled with various textual features to extract relations between genomic anomalies and drug response from MEDLINE abstracts. Our system achieved high precision, recall and F-measure of up to 0.95, 0.86 and 0.90, respectively, on annotated evaluation datasets created in-house and obtained externally from PharmGKB. Additionally, the system extracted information that helps determine the confidence level of extraction to support prioritization of curation. Such a system will enable clinical researchers to explore the use of published markers to stratify patients upfront for ‘best-fit’ therapies and readily generate hypotheses for new clinical trials.

## Introduction

Rapidly evolving molecular profiling technologies have enabled improved detection of alterations in genomic biomarkers that predict response to cancer treatments. This in turn has led to a dramatic rise in the number of studies analyzing the effects of tumor-specific alterations on drug response. With the exception of a few well-established biomarkers such as KRAS in metastatic colorectal cancer [1–3], HER2 in breast cancer [4–6], BRAF in melanoma [7, 8] and EGFR in non-small cell lung cancers [9–11], predictive biomarkers are still not widely adopted in clinical practice due to several challenges. One major issue is that not all genomic alterations are potentially responsive to cancer treatments, i.e. clinically actionable [12, 13]. Actionable genomic alterations help inform individualized treatment plans for cancer patients while minimizing toxicity caused by standard therapy. The percentage of such clinically actionable mutations in tumors is very small and the effect of most genomic alterations with respect to cancer therapies is still unknown. Numerous pre-clinical and clinical studies are being conducted to address this issue and guidelines are being published to classify such genomic alterations based on actionability [14]. Nevertheless, the large volume and complexity of cancer precision medicine literature makes it challenging for busy oncologists and clinical researchers to sort through vast amounts of data and review pertinent information that can inform personalized treatment plans for their patients. Several large scale consortiums such as ClinGen [15], ClinVar [16], My Cancer Genome [17], and CIVic [18] have ongoing efforts to standardize and organize large scale
information linking genomic variants to phenotypic data to drive precision medicine research. UniProt [19], BioMuta [20], OMIM [21], UMD [22], HGVbaseG2P [23], MutDB [24] and dbSNP [25] are few other repositories that house mutations as well as related disease and phenotype information. PharmGKB [26] curates genetic variants and their impact on drug response and diseases. However, all these efforts are based on meticulous manual curation of literature by experts, which is labor intensive and expensive. While expert curated data is highly accurate, the simple task of searching the PubMed database and sorting through thousands of non-relevant papers in order to identify the relevant ones can be extremely time consuming. In a previous study conducted by our group, we generated an expert curated gold-standard corpus of literature on the predictive effect of gene or protein expression of seven biomarkers on response to chemotherapy [12]. During this study, a simple PubMed query to find the co-occurrence of “ERCC1 expression” and platinum-based drugs in the timespan between 01/01/1990 – 12/31/2015 rendered approximately 575 papers out of which only 85 papers were relevant for curation. This shows that even manually identifying the relevant articles for curation is hugely time consuming and expensive as the volume of biomedical articles grows exponentially. Furthermore, there is a lack of resources that researchers can use to obtain and analyze information regarding personalized treatment options based on genomic profiles. Therefore, there is an urgent need to develop an automated approach that can extract relevant context about the effect of genomic alterations on outcomes associated with cancer therapy from literature in order to assist expert curators and clinical researchers.

With the realization of the importance of biomedical text mining, there have been numerous tools developed for various purposes [27, 28]. There have been several text mining tools to extract information from the literature in the pharmacogenomics area, too. Such tools need to first identify mentions of biomedical entities from literature. Currently available tools extract different biological entities [29] such as genes [30], diseases [31], chemicals [32], mutations [33] and species [34]. Additional tools to identify relationships between these entities have been developed. mutation - disease [35, 36] from scientific literature. There are certain tools in pharmacogenomics domain that identifies relationships between drugs and other entities. SNPshot [37] finds binary relation between entities such as mutation-drug, allele-drug and gene-drug using the co-occurrence information of entities along with parse tree and keyword matching. Xu et al. [38] developed a conditional relationship extraction approach to extract drug-gene pairs from MEDLINE abstracts using known drug-gene pairs as prior knowledge. Rance et al. [39] used a co-occurrence based approach to automatically extract mutation-drug relations. Rinaldi et al. [40] modified an existing protein-protein extraction tool to adapt to the extraction of pharmacogenomics relations. Pakhomov et al. [41] proposed to use the positive and negative labels for drug-gene relations from PharmGKB as training data to build a support vector machine classifier to predict drug-gene relationships. Garten et al. [42] developed a text mining tool named Pharmspresso for extracting pharmacogenomics concepts and relationships from full text. Additionally, review articles from Garten et al. [43] and Coulet et al. [44] provide a rich description of the state-of-the-art extraction of pharmacogenomics information from the biomedical literature.

To the best of our knowledge, there is no publicly available text mining tool that extracts entity relationships to indicate the impact of genomic anomalies on cancer therapeutics. This will help biocurators and researchers quickly shortlist relevant precision medicine literature and minimize manual effort. Therefore, our goal was to develop a natural language processing (NLP) based text mining (TM) system, eGARD (extracting Genomic Anomalies association with Response to Drugs), to extract relations between genomic anomalies and drug response from MEDLINE abstracts. This system will in turn support and speed up the process of manual curation. We consider a variety of genomic anomalies such as substitutions, duplications, insertions, deletions, gene copy number variations, structural variants as well as gene and protein expression changes. eGARD accounts for a variety of ways that relationships between anomalies and drug responses are described in text, unlike majority of the existing TM systems that mostly use co-occurrence of entities. We make use of the syntactic nature of the sentences coupled with various textual features for extraction. To assist with such extraction of information, we have extended and repurposed multiple in-house and public text mining systems including DiMeX [45], miRiaD [46] and PubTator [29]. Moreover, we extract additional details about the study to help experts determine the quality of evidence presented in the study. We evaluated our system on different annotated datasets; both created in-house and obtained externally (PharmGKB). We achieved precision, recall and F-measure of up to 0.95, 0.86 and 0.90, respectively.

eGARD was applied on roughly 36,000 PubMed abstracts, that were retrieved for 50 genes and 42 cancer drugs including cell cycle inhibitors, kinase inhibitors and antibody treatments, illustrating the scalability of the system. It produces output in JSON format to facilitate data exchange and integration of text mining results. The JSON representation can enable the use of existing curation tools for expert validation and ranking. All extracted results are stored in a database and are available for curators and clinical researchers. The extracted information relating genomic anomalies to drug responses will enable researchers to readily generate hypotheses for new precision medicine based clinical trials.

## Materials and methods

In this section, we describe how eGARD extracts the association between a genomic anomaly and drug response. It begins with a discussion of the task itself and the type of sentence structures we focus on. The latter discussion describes the rules we have developed to extract relationships from sentences. Following this discussion, we introduce the details of the actual method including the processing of text, entity recognition and typing and pattern matching.

### Defining the task

The goal of this work is to identify the association between genomic anomaly and drug responses from text (currently MEDLINE abstracts). Example 1 provides a sample sentence that captures the information eGARD is designed to extract. The connection stated in this sentence is expressed by “was significantly associated”.

> Example 1: “<u>Low expression of Bax</u> was significantly associated with <u>poor survival</u> of patients with metastatic or recurrent *gastric cancer* treated with *FOLFOX* chemotherapy.” (PMID: 20503071)

The genomic anomaly in Example 1 is expressed as “low expression of Bax”. eGARD not only considers anomalies that are different gene or protein expression levels but also variants. The latter includes substitutions, duplications, insertions, deletions, gene copy number variations and structural variants. We will refer to all of them as *Anomaly entity* in this article. eGARD captures phrases like “Over-expression of ERCC1”, “Patients with high RRM1”, “PAX4 variant rs6467136”, “IL28B polymorphisms” that express anomalies.

In order to capture the effects of genomic anomalies on responses to treatment, we are interested broadly in concepts such as survival or response rate after drug treatment, change in sensitivity to drugs, and outcome of treatment which includes overall survival and progression free survival. While there could be more terms that represent the effects, these concepts are of key concern to precision oncology, explaining their prevalence in the associated literature. We will refer to them as *RO entity* (RO stands for Response or Outcome) in this article. We treat phrases such as “significantly poorer response”, “survival rate”, and “increased drug resistance” as RO entities. A formal definition of how RO entities are detected is described later in this article in the subsection “Entity recognition”. Example 1 associates RO entity term “poor survival” with the Bax’s low expression level. Clearly, the information about the response/outcome is incomplete without the extraction of drug/treatment (“Folfox chemotherapy” in Example 1) and the disease for which this treatment is being used (captured by the phrase “metastatic or recurrent gastric cancer”). eGARD attempts to capture these four components from text, viz., the anomaly, the response/outcome, the drug and the disease. Here, all four are mentioned in the same sentence. However, this is not always the case and sometimes the disease and the drug has to be inferred from the rest of the abstract.

Finally, in addition to the extraction of the four components from sentence shown in Example 1, eGARD also extracts the information that the low expression level of Bax has a *negative impact*. In general, for an association between an anomaly and a RO entity, eGARD attempts to classify whether the impact is one of *benefit, lack of benefit* or *not*-*assessable*.

## Different sentence types

The association of genomic anomaly with drug responses for certain drugs and diseases are expressed in text in different ways. Our approach is to detect certain lexico-syntactic dependency structures in sentences to extract the relations between an Anomaly entity and a RO entity. The drugs and disease will be extracted from context, whether from the sentence or the rest of the abstract. Based on our observation, we have identified several types of sentence structures that convey the relationships that we are interested in. These different sentence structures together with our rules for extraction are described below. Our method for identifying the drug and disease entities is discussed following the description of different sentence types.

### Association type

There are sentences that mention the relationship between genomic anomaly and drug responses in the form of an association. Usually there is a “trigger” word that indicates the association relation between an Anomaly entity and a RO entity. In Example 1 the trigger word is the “associate” itself and in Example 2 below, the trigger word is “correlate”. There are several other trigger words (such as “relationship”, “contribute”, “(play a) role”) that we use for this sentence structure, with several textual variations for each trigger.

> Example 2: “<u>The MGMT expression</u> was inversely correlated with <u>response to</u> <u>temozolomide</u>.” (PMID: 20130512)

As with Examples 1 and 2, the Anomaly entity and the RO entity serve as syntactic arguments of the trigger word. The syntactic relations are of subject and object of the trigger, and the trigger word is a verb. Our rules for association type sentence structure look for the syntactic pattern: <*anomaly entity*> *“trigger word” <RO entity*>. Our system first identifies sentences which contain the trigger word. In such cases we check if the requisite syntactic arguments of the trigger are an Anomaly entity and a RO entity. In this case, the pair is extracted.

As mentioned earlier, eGARD attempts to find the impact type (one of *benefit*, lack of benefit or *nonassessable*) from the extracted associations. In association type sentences, sometimes the trigger verb is modified by an adverb which indicates the impact type. Example 2 is one such case where the adverb “inversely” indicates that MGMT expression will inversely impact the temozolomide response, from which we infer that high expression levels of MGMT incurs *lack of benefit* from temozolomide. Other than “inversely”, we use words “positively” and “negatively”, too. If we cannot determine impact type from the adverb modifier, we look into the Anomaly entity and RO entity to determine the impact type. From the Anomaly entity, we detect information such as expression levels by looking at adjective modifiers “low”, “high”, “elevated”, “overexpressed” etc. or specific constructs such as gene name followed by words “positive”, “negative”, “deficiency” etc. From the RO entity, we try to detect the impact type by the presence of certain adjectives modifying the drug responses, such as “poor”, “better”, “greater”, “worse” etc. For instance, in Example 1, we detect the expression level of Bax as “low” from the Anomaly entity and drug response as “poor” from the RO entity. Combining both, we infer that low expression of Bax incur *lack of benefit* for drug response.

### Comparison type

This type of sentence also expresses an association relation between Anomaly entity and RO entity except that the relation is expressed with a comparison. These types of sentences are very common in biomedical literature, especially in the pharmacogenomics domain. Unlike the association sentence type, there are three entities involved in the comparison sentence type: an observed entity and two compared entities. We are interested in comparison sentences where an observed entity is an RO entity and the two compared entities are related to the Anomaly entity (as in Example 3 below) or vice-versa.

> Example 3: “<u>ERCC1-negative patients</u> had better <u>PFS (P = 0016) and OS (P = 0030)</u> compared with <u>positive patients</u>.” (PMID:25107571)

In Example 3, the observed entity is indicated by the phrase “better PFS (P = 0016) and OS (P = 0030)”, and the two compared entities are “ERCC1-negative patients” and “positive patients”, respectively. Therefore, this comparison marks an association between ERCC1 expression level and survival of patients.

To recognize and extract information from comparison sentences, we look for two clues. First, we consider the trigger to be a comparative adjective (with the part of speech of JJR in the parse structure) such as higher, lower, greater, better etc. (“better” in Example 3). Then, we look for phrases such as “compared with”, “in comparison to”, “compared to”, “than” and “versus” which often separates the two compared entities.

The comparative adjective is connected to the observed entity in one of two ways. It can modify the observed entity which appears as noun to its right as in Example 3. The syntactic pattern that corresponds to Example 3 is: <*Anomaly entity*> *“trigger” <RO entity*> *comparison_phrase <Anomaly entity*>. Alternatively, the comparative adjective can appear as the head predicate of a sentence as in Example 4 and in these cases, its subject (TS and TP mRNA levels in Example 4) is extracted as the observed entity. The syntactic pattern that corresponds to Example 4 is: <*Anomaly entity*> *in* <*RO entity*> *“trigger” comparison_phrase* <*RO entity*>. Note that in either case, one of the two compared entities appears immediately after the comparison phrase (“compared with”, “compared to”, “than” etc.). The other argument will be either the subject when the JJR modifies a noun or as an adjunct of the subject when it appears as the head predicate (Example 4).

> Example 4: “<u>TS and TP mRNA levels</u> in the <u>patients with complete response, partial response or stable disease (n = 34)</u> were significantly lower compared to those in <u>the patients with progressive disease (n = 11)</u> (p = 0017 and p = 004, respectively).” (PMID: 22783377)

There are several other variations in the comparison structures including cases where only one of the compared entities is explicitly mentioned in the sentence. In such cases, the implicit compared entity is usually implied from the context. Consider the sentence in Example 5 where one of the compared entities is missing in the sentence, but implicitly referred to as groups with high level of TS.

> Example 5: “However, <u>the group with low level of TS</u> had a longer <u>DFS</u> (144 mo versus 83 mo, P = 0017).” (PMID: 17854149)

The impact type in comparison sentences are detected by combining information presented in the trigger (the comparative adjective) and in both Anomaly and RO entities. In the case where the observed entity is the RO Entity (as in Example 3), we detect the expression levels or specific mutations from the Anomaly entity and the impact type from the trigger. Additionally, we recognize the type of the RO entity so that we can correctly qualify the impact type. Let’s consider Example 3 to illustrate our approach. The Anomaly entity (“ERCC1-negative patients”) specifies low expression of ERCC1, the trigger “better” specifies a positive correlation and finally, the RO entity (“PFS (P = 0016) and OS (P = 0030)”) specifies the outcome as survival type (PFS and OS stands for progression-free and overall survival, respectively, determined by the use of an acronym detector).

Combining all of them, we infer that low expression ERCC1 incurs *benefit* for the drug response. However, if the observed entity is the Anomaly entity (as in Example 4), the trigger indicates the expression levels and the first compared entity (which is a RO entity) hints about the impact type. In Example 4, the trigger “lower” represents the expression levels for Anomaly entity (“TS and TP mRNA levels”) and the RO entity (“patients with complete response, partial response or stable disease”) represents response of patients. Combining both, we infer that low levels of TS and TP incur *benefit* for the drug response.

### Sensitization type

Words like “resist” or “sensitive” can appear in their noun form as RO terms in the association type sentences as in Example 6 for instance.

> Example 6: “These results indicate that <u>enhanced MGMT expression</u> contributes to <u>TMZ</u> <u>resistance</u> in MGMT-positive GSCs.” (PMID:23958055)

However, the verb forms of these words are also often used in sentences that connect them with Anomaly entities. These verbs appear as the head predicate of the clause with the Anomaly entities as their subjects. Our rules for extraction look for the following syntactic pattern: <*anomaly entity*> *sensitizes* <*disease (cells)*> *to* <*drug/treatment*> as shown in Example 7. That is, the trigger appears as verb (VBN) with the Anomaly entity appears as its subject and the disease cells as a direct object. The drug or treatment phrase appears as a prepositional phrase modifying the trigger with the preposition “to”.

> Example 7: “<u>ATM deficiency</u> sensitizes mantle cell lymphoma cells to <u>poly (ADP-ribose)</u> <u>polymerase-1 inhibitors</u>.” (PMID:20124459)

Sometimes, the disease cells that are sensitized are not mentioned in the sentence. In such cases, as discussed later, the disease is inferred from the context.

Detection of impact type from sensitization type sentences is fairly straight-forward. In the syntactic pattern mentioned above, as the trigger “sensitize” suggests that the Anomaly entity causes the disease cells to respond to the drug, the Anomaly entity always incurs *benefit* for the drug response. Therefore, it is sufficient to look only at the Anomaly entity involved and determine the expression levels or specific mutations that cause the *benefit*. For example, the Anomaly entity (“ATM deficiency”) refers to low levels of ATM. Thus, low expression of ATM incurs *benefit* for the drug response.

### Marker type

In this type of sentence structures, we consider cases where the genomic anomalies are stated to be *markers* for drug responses. These relationships are triggered by words such as “predictor”, “biomarker”, “marker”, “indicator” etc. For instance, the sentence in Example 8 is of this type of sentences with trigger word “predictor”.

> Example 8: “Multivariate analysis showed that <u>low expression of ERCC1</u> was an independent predictor for <u>prolonged survival</u> (HR, 0120; 95% CI, 0016-0934, P = 0043).” (PMID:18594541)

A key requirement for these sentences is the presence of an “is-a” verb group. We have adopted the approach mentioned in miRiaD [46], which uses “is a”, “are”, “acts as”, “functions as”, “serves as” and appositives as triggers for the “is-a” relation. The Anomaly entity is found as the subject of “is-a” and the “marker” trigger is its object. The RO entity is found as the noun modifier for the trigger often linked with the preposition “for”. Using this rule, from Example 8, we can extract “low expression of ERCC1” as the Anomaly entity (from which the gene and anomaly are obtained), “prolonged survival” as the RO entity.

Given the nature of copular sentences, we can detect variations of this type of sentences where the trigger might be an adjective (JJ) instead of noun, such as “predictive”, and “indicative”. Example 9 presents one such sentence where the trigger is the adjective “predictive”.

> Example 9: “Furthermore, <u>concomitant low expression levels of ERCC1, RRM1, and RRM2</u> <u>and the high expression level of BRCA1</u> were predictive of a <u>better outcome</u> (P = 0014).” (PMID:25227663)

Impact type detection in these cases is identical to impact type detection in association type, where we combine information from both the Anomaly and RO entity and infer the impact type. For instance, applying the rules mentioned earlier, Example 8 will yield the information that low expression of ERCC1 *benefits* the drug response (survival in this case).

### Statistical type

There are sentences that do not explicitly indicate the association between genomic anomaly and drug responses. However, these sentences seem to present a quantitative value of an RO entity for multiple and often contrastive Anomaly entities. When an expression of a P-value is given following the quantities, we hypothesize that there is an implicit comparison to draw an association (See Example 10 below). We do not detect the impact type since it can be inferred only by comparing the numbers, and knowing how to interpret the numbers.

> Example 10: “Among cohort 2, <u>the response rates</u> of <u>patients with low ERCC1 and high</u> <u>ERCC1 expressions</u> were 45.5% and 20.0% respectively (P = 0.361).” (PMID:23358102)

Here the outcome is mentioned in terms of response rates and is connected to ERCC1 levels.

### Architecture of the system

The schematic diagram in Fig 1 shows the overall workflow of our system. It starts with text preprocessing of MEDLINE abstracts and syntactic processing of the text. The parsed phrases are then typed to different categories, which is followed by the extraction of association of genomic anomaly and drug response. We additionally extract other information that helps determine the confidence and evidence level of extraction. Finally, the results are disseminated in convenient format such as JSON.

**Fig 1:**
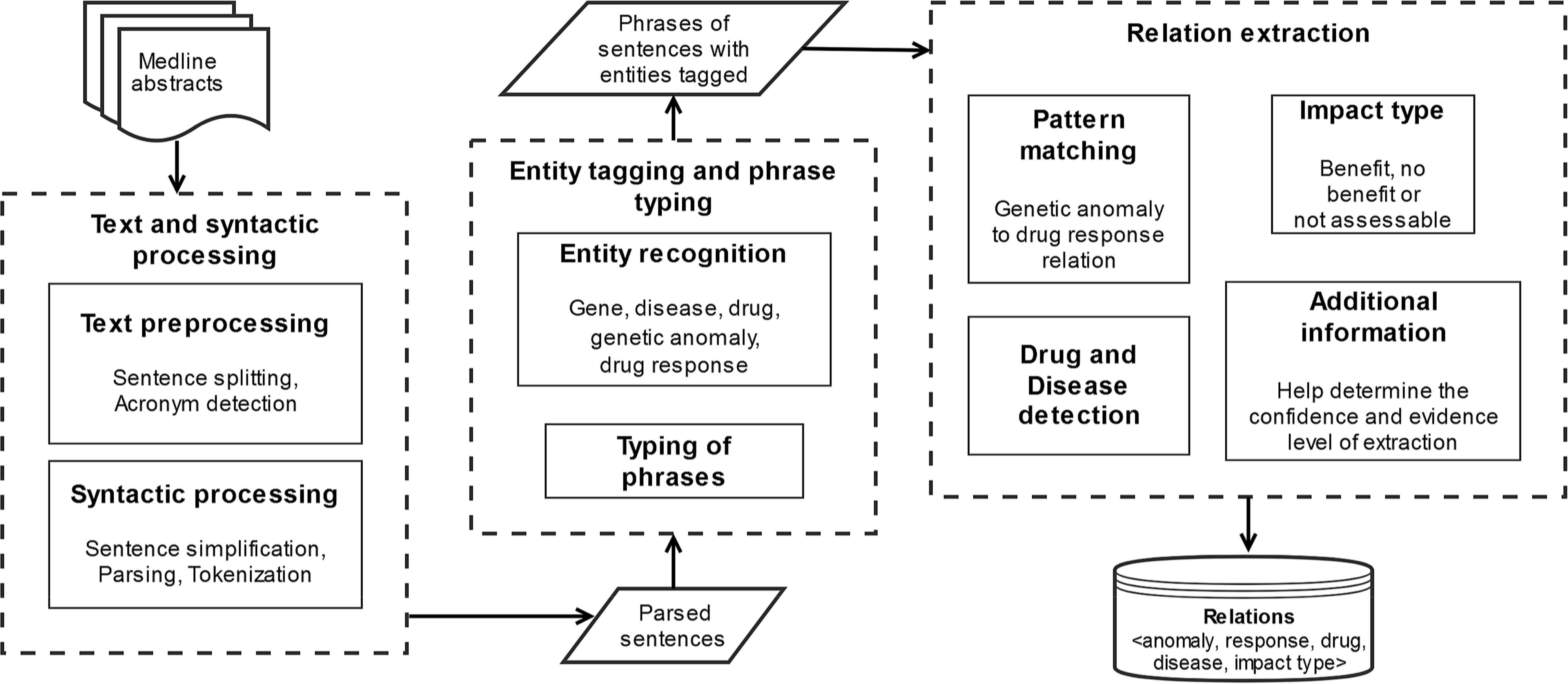
Schematic diagram of the developed system.

### Text preprocessing

Our system takes PMIDs as input and retrieves the title, abstract and MeSH terms from MEDLINE repository. An in-house sentence splitter was used to split the abstracts into individual sentences. We used an acronym detector [47] to detect possible abbreviations. The abbreviated forms of different terms and entities assisted the entity detection step. For instance, in the text excerpt “The median disease-free survival (DFS) time was 10.2 mo in the patients.” (PMID: 17854149), the term “disease-free survival” is a RO entity and the acronym detects the abbreviated form for it as “DFS”. Thus, we treated DFS as a RO entity as well throughout the abstract.

### Syntactic processing

We employed sentence simplification by Yifan et al. [48] in some cases to simplify complex sentence syntaxes into simpler forms. This facilitates the extraction of relations with simpler and more uniform patterns instead of applying complex patterns. For instance, consider the sentence in Example 11.

> Example 11: “High MDR1 and ERCC1 gene expressions are associated with inferior outcome after cisplatin-based adjuvant chemotherapy for locally advanced bladder cancer”. (PMID: 20689757)

The simplification step will render these simplified sentences: “High MDR1 gene expressions are associated with inferior outcome” and “High ERCC1 gene expressions are associated with inferior outcome”. Thus, applying one simple pattern, we can extract both the relations without explicitly handling conjunctions in the pattern.

We used BioNex [49] for tokenization and parsing of sentences into chunks of base noun phrases (NP) and base verb groups (VG). When consecutive base NPs are connected via prepositions, conjunctions or punctuation marks, we merged them to form larger NPs. Similarly, consecutive base VGs were merged into larger VGs as well. For example, the BioNex system will parse the sentence is Example 11 to the bases phrases as: *NP(High MDR1), NP(ERCC1 gene expressions)*, *VG(are associated)*, *NP(inferior outcome)*, *NP(cisplatin-based adjuvant chemotherapy)* and *NP(advanced bladder cancer)*. The merged phrases will be *NP(High MDR1 and ERCC1 gene expressions)*, *VG(are associated)* and *NP(inferior outcome after cisplatin-based adjuvant chemotherapy for locally advanced bladder cancer)*.

### Entity recognition

To detect diseases, we used annotation from Pubtator [29], which is a publicly available tool that assists biocuration by tagging various biological entities. We downloaded and used the pre-computed annotations from Pubtator. The disease mentions are normalized to MEDIC IDs, which again we normalized to DOID in Disease Ontology (DO) [50] database. We noticed some mistakes in disease tagging in Pubtator, such as *AR* being commonly tagged as disease although its full form also mentioned in the abstract is the gene *Androgen Receptor*. The system was able to automatically rectify this type of problems using the acronym detector. The acronym detector detects AR as a short form of Androgen Receptor, which in turn is detected as a Gene. By looking at the full form detected by acronym detector, the system was able to discard AR a disease and consider it as a gene mention. For gene mentions, we used both Pubtator and an in-house gene mention detector pGenN [51]. The gene mentions were normalized to EntrezIDs. For drug detection, we used Pubtator annotation, too.

Genomic anomalies can be either mutations or change in gene expression levels. Although they are biologically different, they both have the potential to impact the drug responses of a therapy. Therefore, we treat them similarly in the context of our work, grouping them together under Anomaly entity. Mutations were detected using our previously built tool used in the system DiMeX [45], which also provides mutation to gene associations. Additionally, we developed a separate system to detect other kinds of genetic variations such as mentions of copy number variations (CNV) and gene amplifications and loss/gain of function mutations.

We developed our own module to detect gene expression level mentions as genomic anomalies. Usually expression terms appear along with the corresponding gene name in the same noun phrase, with the head word of a NP indicating an expression (e.g., expression, overexpression, inhibition, deficiency, levels etc.). Sometimes the expression terms are connected to the gene name via the “of’ preposition (e.g “high expression of TS”). We also detect NPs as expression entities if they are headed by gene names and modified by level indicators like “low” or “high” (e.g., like “low TS”, where TS is a gene name). Additionally, phrases that indicate expression levels with numeric values along with a gene name such as “TS <= 7.5×10 (-3)” are also detected as expression entities.

The RO entities are detected by NPs headed by words that indicate response or outcome. Based on our observations, we have identified several of such words, such as “survival”, “prognosis”, “outcome”, “response”, “efficacy” etc. because the NPs headed by these words represent an outcome when the drug(s) are administered. Thus, these NPs represent our definition of drug responses. Some examples of such detected NPs are “longer survival”, “higher response”, “poor efficacy”, “overall increased response” etc. We have also used a list of fixed phrases (and their corresponding acronyms, if applicable) that indicate RO entities. Some examples of such occurrences are “progression free survival”, “PFS”, “overall survival”, ‘OR”, “objective response rate” etc.

Table 1 summarizes the tools that were used for various entity detections.

**Table 1.** List of tools used for different entity detections.

| Entity | Tool |
| --- | --- |
| Gene | Pubtator + PGenN |
| Disease | Pubtator |
| Drug | Pubtator |
| Mutation | DiMeX |
| Copy Number Variation | eGARD |
| Gene expression | eGARD |
| Drug response | eGARD |

### Typing of phrases

Once the noun phrases (NP) are obtained from the parser, we categorize the NPs depending on the entities that appear within the NP, such as gene, disease, mutation etc. We named this step as typing of phrases, as the NPs are assigned an entity type. The entity type is assigned based on occurrences of certain entities or keywords at the head of the NP. We took the rightmost word of an NP as the head. If multiple base NPs were merged together due to prepositional phrases attachments to form one NP, we consider the head of the leftmost constituent NP to be the head of the entire merged NP. For instance, consider the sentence in Example 12. The first NP (“High expression of thymidylate synthase”) is of type <gene-expression> because the head word (“expression”) of the first constituent NP (“High expression”) refers to expression type. Likewise, the second NP (“the drug resistance of gastric carcinoma to high dose 5-fluorouracil-based systemic chemotherapy”) is taken to be of type <drug-response> as the head word of the first constituent NP is “resistance”.

> Example 12: “<u>High expression of thymidylate synthase</u> is associated with <u>the drug resistance</u> <u>of gastric carcinoma to high dose 5-fluorouracil-based systemic chemotherapy.</u>” (PMID: 9576280)
>
> However, in certain cases, the head of the leftmost constituent NP was not conclusive for any type such as when the head word is patient or group. In such cases we consider the head of the next constituent NP that modifies it to determine the type of the NP. For example, in the text excerpt “patients with low ERCC1 expression showed a significantly higher rate of good tumor response” (PMID: 25674147), the type of the NP “patients with low ERCC1 expression” was determined to be <gene-expression>. Using the same approach, NPs were typed to other entities, namely <disease>, <drug>, <mutation>, <gene> etc.

### Pattern matching

In the discussion of the different sentence types above, we already outlined some of the syntactic patterns that we used to match against text. In this section, we will formally define the patterns and discuss the matching process with an example. Let’s reconsider the syntactic pattern for sensitization pattern that we introduced earlier: <*anomaly entity*> *sensitizes* <*disease (cells)*> *to <drug/treatment*>. This pattern can be formally broken down and written as:

sensitize_VG *has*_*subj* Anomaly_NP;
sensitize_VG *has*_*obj Disease*_ cell_NP;
sensitize_VG *nmod*_*to* Drug_NP

The common element that binds the entire pattern together is a verb group (VG) headed by the word “sensitize”. So, to match the pattern in text, we first searched for a (possibly merged) VG with head “sensitize”. As we already have the types of NPs (from the typing of phrases step), the next step is to ensure that the *has*_*subj* constraint is met by finding an NP of type <Anomaly> that is a subject to the detected sensitize_VG. This is done by checking if an NP of type <Anomaly> appears immediate left of the sensitize_VG when this verb group is in active form (detected by the parser). Similarly, the *has*_*obj* constraint is ensured by looking for a NP of type <disease_cell> to the immediate right of sensitize_VG. Finally, the *nmod*_*to* constraint is ensured by searching a NP of type <Drug> that is a modifier for the sensitize_VG and is connected as a prepositional phrase via “to”.

Similarly, the syntactic pattern for one of the association type sentences, <*anomaly entity*> *associate* <*RO entity*>, can be formally written as follows and the pattern is matched in text using the same technique.

associate_VG *has*_*subj* Anomaly_NP;
associate_VG *has*_*obj* RO_NP

All the patterns that are used in this work are matched using this same approach. List of all patterns for all sentence types are available as supporting information (S1 File).

### Finding the disease and drug

In addition to detecting association between genomic anomaly and drug responses, eGARD also records the associated drug and disease. The drug and the disease may co-occur in the same sentence that mentions the association between the Anomaly entity and the RO entity. In such cases, if the drug and/or disease is found in the noun phrases that are extracted for Anomaly or RO entities by the patterns, then they are extracted as well. However, quite often, the drug and/or disease are not mentioned in the same sentence but must be inferred from the context. Based on our observations, we have applied certain heuristic techniques to locate the drug(s) and disease(s) from context in such cases (i.e., where they cannot be extracted from the same sentence).

<u>Drug detection:</u> To detect the drug that corresponds to the drug response, we looked for some simple patterns at certain rhetorical zones in the abstract in the following order: title, method sentences, patient context (PC) sentence, conclusion sentence(s) and introduction sentence(s). A PC is a sentence in the abstract which mentions information about the patients involved in the study. To detect PC sentence and split the abstract into five rhetorical zones (Title, Introduction, Method, Result, and Conclusion); we adopted the implementation mentioned in DiMeX [45]. If we denote the mention of drugs as *drugname*, some of the patterns that we used are but not limited to “treatment with *drugname*”, “patients treated with *drugname*”, “patients receiving *drugname*”, “*drugname* therapy”, “efficacy of *drugname*” etc. The intuition behind these patterns was to identify the drugs that were used to treat patients, rather than just looking for co-occurrence of drugs. The full list of patterns used is available as supporting information (S1 File).

<u>Disease detection:</u> We consider the disease mentioned in the Patient Context (PC) sentence to be the central disease of the study, as in DiMeX [45]. Thus, we look for the disease in a PC sentence, provided such a sentence occurs before the current sentence. If the disease is not found in a PC sentence, we look for the central disease at other rhetorical zones in the abstract in the following order: title, conclusion sentence(s) and introduction sentence(s).

### Additional information extraction

In addition to the extraction of the association of genomic anomaly and drug responses, we also extracted additional information that we believe would be helpful for a curator or a researcher to easily distinguish information extracted from a patient vs. cell line study or prospective vs retrospective study. This may be useful in easily determining and summarizing the level of evidence associated with a predictive biomarker. Firstly, we extract information related to patients in the study, such as size of the experimental patient population and control, the race or nationality, etc. For detecting patient or control size, we used regular expressions. To detect the race/nationality of patients, we used a pre-compiled list country names, their adjectival and demonym forms. This information is assumed to indicate whether the investigation involved patients (as compared to those on cell lines or models) and hence can be used to prioritize curation or rank the importance of the extracted conclusion. We also look for NPs of type <cell> or <cell-lines> (given by the typing of phrases step) and tag the abstract as being related to cell type study instead of patient study. Additionally, we look for the presence of certain information (in the form of words or phrases) in the abstract that provide valuable insight for a curator or a researcher to filter or rank information. These information include but not limited to: retrospective or prospective study, clinical trial phases (I or II or III or IV), in-vivo or in-vitro or ex-vivo, clinical trial IDs and meta-analysis. We match these phrases or their minor variations against abstract text. Finally, we check if the publication is a review article by examining the MeSH terms.

### Evaluation setup

We evaluated our system on different datasets to evaluate how well eGARD performs over a range of text. Our data sets included two sets of abstracts that were annotated in-house. We also evaluated the system on a set of abstracts based on the PharmGKB [26] data. For evaluation, we counted true positives (TP), false positives (FP), and false negatives (FN), true negatives(TN) and used the standard information retrieval metrics of Precision (P), Recall (R), F-measure (F), and True Negative Rate (TNR) for performance evaluation, where P = TP/(TP+FP), R = TP/(TP+FN), F = 2PR/(P+R) and TNR = TN/(TN+FP).

### In-house annotated sets

We used a set of abstracts from which the information regarding the association of genomic anomaly on drug response had already been annotated during a previous curation work [12]. The curated information had been recorded along with the articles’ PMID and sentences in the abstract that conveyed the information. This curation was done by one of the authors, a domain expert who did not participate in the design and implementation of the eGARD system. The same author was able to quickly convert the information into annotated dataset that could be used for evaluation. We called this set InHouseSet1. It includes 100 abstracts, where each abstract is annotated with information containing a gene, the type of anomaly (specific mutations, high or low expression levels etc.), the drug and the disease. Furthermore, the type of impact (benefit, lack of benefit or not assessable) of the anomaly on the response is also recorded. The abstracts in InHouseSet1 pertain to seven gene-drug combinations. The list of these seven gene-drug combinations is available as supporting information (S2 File).

We consider the annotation to be correct only if all 4 components that are extracted by the system matched with those in the annotation. The results are reported in terms of precision, recall and F-measure. Since the impact type can be found in the annotation only for the true positives for the extraction of the 4-tuple, we separated the evaluation of the impact type from the extraction of the 4tuple. Hence we considered the impact type annotations only when the system correctly extracts the 4-tuple and did not consider the 4-tuples incorrectly extracted by the system, which obviously will not have any associated impact type information in the annotation.

We also considered a second in-house annotated dataset. A PubMed search for a biomarker gene and drug combination often returns hundreds of abstracts, where many might not contain information on the genomic anomalies’ association with drug response. For example, we reported in the Introduction that only 85 abstracts of the 575 abstracts returned from such a search was relevant. In that sense, InHouseSet1 is different from a typical search result because nearly all of them were relevant for curation. Thus, this set is not appropriate to evaluate the system’s ability to reject an abstract as irrelevant and thereby save precious curation time and effort. Thus, we developed another dataset, called InHouseSet2, which was annotated by the same annotator. The only difference is that the 100 abstracts in InHouseSet2 were chosen randomly from the results of a PubMed search on the same seven gene-drug combinations that appeared in the InHouseSet1 data. In contrast with InHouseSet1, only 38 of the 100 abstracts in InHouseSet2 contained relevant information.

Since the primary goal of developing and using InHouse2 for evaluation was to consider the ability to reject irrelevant abstracts, determination of true negatives is important. Thus, we focus on the metric of True Negative Rate (TNR), also known as specificity and defined as TNR= TN/(TN+FP), when using InHouseSet2. We also provide the precision and recall results, although it must be noted that the number of positive instances are much smaller and hence the precision and recall results are less reliable in contrast to the use of InHouseSet1. Both InHouseSet1 and InHouseSet2 are available as supporting information (S1 Dataset and S2 Dataset, respectively).

### PharmGKB dataset

To evaluate how well our system’s performance generalizes, we considered a dataset that was not annotated in-house. For this purpose, we used the data from the PharmGKB project. PharmGKB has a variant annotation dataset containing manually curated associations in which the variant affects a drug dose, drug response or drug metabolism. We took the PharmGKB variant annotation dataset and tailored it to use it to evaluate our system’s ability to extract the impact of variants on drug responses. This evaluation provides a chance to examine how much of manually curated information in an existing database our system can reproduce. As our work is concerned about drug responses, we filtered out effects of variants on drug dose and metabolism, keeping only the drug responses. In this filtered version, we searched for annotations that are concerned with a list of FDA-approved drugs for 13 genes in cancer study. This list of drugs (and their variations in names), which is available as supporting information (S3 File), was provided to us by the annotator. This yielded a set of 46 articles. From the rest of the filtered annotation set, we randomly picked 54 articles to make the entire PharmGKB evaluation set containing 100 articles. As our system is concerned with abstracts only, we did not include the annotations where the information is found in the full-length text but not the abstracts. The above filtering process yielded set of 76 annotated associations from 46 abstracts (FDA) and 65 annotated associations from 54 abstracts (others), making the entire set of 100 abstracts with 141 annotated associations.

We used recall value as evaluation metric for this set. However, because all of the annotations are positive and we cannot calculate false positives, we cannot assess the precision for this set. Also, PharmGKB annotation does not contain disease information and hence it was not included in the evaluation. Additionally, since the PharmGKB was not annotated for impact information, this aspect also cannot be evaluated by this dataset. The PharmGKB dataset was obtained via acquiring a license through https://www.pharmgkb.org/. Since the annotation was not developed in-house, we only list the PMIDs for the 100 abstracts as supporting information (S3 Dataset).

## Results and discussion

We first report the results of eGARD when it was evaluated on the InHouseset1 data set. Table 2 records the precision and recall for the extraction of the four components: the anomaly, the response, the drug and the disease. We consider an extraction to be correct only if all four components match the corresponding components in the annotation. We have achieved F-measure of 0.90 with a precision of 0.95 and recall of 0.86. In depth analysis of the mistakes revealed mainly three types of errors. First was due to the complexity of sentences, as in the sentence “Gene expression levels of APTX, BRCA1 and ERCC1 were significantly lower in irinotecan-sensitive gastric cancer samples than those irinotecan-resistant samples (P<0.001 for all genes), while ISG15 (P=0.047) and Topo1 (P=0.002) were significantly higher.” (PMID:23517622). While eGARD was able to extract the relationship between APTX, BRCA1 and ERCC1 expressions with irinotecan sensitivity, it failed to capture the relation between ISG15 and Topo1 with the drug response. We had previously mentioned that the drug and disease might not be mentioned in the same sentence and that eGARD attempts to identify them from context in these cases. A few errors were due to the extraction from context.

Finally, while the patterns in eGARD gave us a recall of 0.86, there were a few FNs due to missing patterns. For example, “Class III beta-tubulin overexpression is a prominent mechanism of paclitaxel resistance in ovarian cancer patients” (PMID:15671559) was missed because of the lack of a trigger.

**Table 2.** Performance of our system in finding the association of a genomic anomaly with drug responses (<anomaly, response, drug, disease>) in InHouseSet1.

|  | TP | FP | FN | Precision | Recall | F-measure |
| --- | --- | --- | --- | --- | --- | --- |
| InHouseSet1 | 128 | 7 | 21 | 0.95 | 0.86 | 0.90 |

We additionally evaluated our system’s detection of impact type (one of three: *benefit*, *lack of benefit* or *not assessable*). For this study, as explained earlier, only the 128 TP cases can be considered. eGARD obtains an 78% accuracy in detecting impact type among these cases. Majority of the mistakes were due to the fact that impact type was mentioned not in the same sentence from which the impact information was extracted. For example, from the sentence: “TUBB3, TS and TP expressions could predict the response of advanced gastric cancer patients receiving capecitabine-based and paclitaxel-based chemotherapy” (PMID: 23358102), the expression levels of the genes and the response quality (either better or worse) are not known. However, a human reader can infer the impact type from earlier sentences where the response rate was quantified for different expression levels. There are few cases where our system was not able to identify the impact type even if it could be inferred from the same sentence. For instance, the system successfully detected the association from the sentence “Increased activity or expression of MRP1, Bcl-xl, TS, and E-cad appear to be involved in the MDR mechanism of BEL-7402/5-FU” (PMID:23167424), but failed to determine that “MDR mechanism” is not beneficial for the treatment (MDR is an acronym for multidrug resistance).

As mentioned earlier, we have developed another dataset, InHouseSet2, to evaluate eGARD’s ability to differentiate between relevant and non-relevant abstracts. Therefore, evaluation of this dataset was performed at the abstract level, meaning each abstract was considered either relevant or non-relevant, as opposed to inHouseSet1 where individual combinations of the four components were annotated. The result of this evaluation is presented in Table 3. We used the true negative rate (TNR) value to represent how well our system rejects non-relevant abstracts. There were only two cases in this 100 abstract where eGARD predicted an abstract to be relevant when in fact, it was not according to the annotation (false positives), thus yielding a TNR value of 0.97. This is an encouraging result because it suggests that eGARD can considerably save curation time and effort. Although we also present the precision and recall in Table 3, we would like to note that the number of positive abstracts in this set is small.

**Table 3.** Performance of our system in separating non-relevant abstract from relevant abstracts in InHouseSet2.

|  | **TP** | **FP** | **FN** | **TN** | **Precision** | **Recall** | **F-measure** | **TNR** |
| --- | --- | --- | --- | --- | --- | --- | --- | --- |
| **InHouseSet2** | 28 | 2 | 10 | 60 | 0.93 | 0.74 | 0.82 | 0.97 |

The evaluation results for PharmGKB set is presented in Table 4. Please note that we are only able to assess the recall value for this set, as it only contains positive annotations. The overall recall on the 100 abstracts is 0.77. Table 4 shows that similar recall scores are obtained for the two subsets: abstracts based on 13 FDA approved cancer drugs, and abstracts for the other drugs (which covered mostly non-cancer diseases).

**Table 4.** Performance of our system in finding the association between genomic anomaly and drug responses in PharmGKB dataset.

|  | **TP** | **FN** | **Recall** |
| --- | --- | --- | --- |
| **PharmGKB** (FDA drug set, 46 articles) | 59 | 17 | 0.78 |
| **PharmGKB** (other set, 54 articles) | 50 | 15 | 0.77 |
| **PharmGKB** (combined) | 109 | 32 | 0.77 |

## Large scale run

A vast amount of information regarding the association of genomic anomalies and drug responses is already available in published literature. To illustrate the robustness and how well it scales up, eGARD was applied on a large set of PubMed abstracts. We searched for combinations of 50 genes and associated cancer drugs in PubMed that yielded 35,677 abstracts that contains abstract text. Table 5 lists some of the key characteristics from applying eGARD on those abstracts. It can be noted that only 7,309 abstracts (20%) were deemed relevant by our system. From those 7,309 abstracts, our system detected 20,282 associations between a genomic anomaly and a drug response.

**Table 5.** **Characteristics of the extracted results from the large scale run. The abstracts were retrieved for 50 <gene, cancer drug> pairs from PubMed search**.

| Characteristics | Counts |
| --- | --- |
| Abstracts | 35,677 |
| Abstracts with at least one associations | 7,309 |
| Total anomaly-drug response associations | 20,282 |

## Conclusion

In this work, we have described a text mining system eGARD that addresses the needs and challenges of automatic extraction of information regarding genomic anomalies and their impact on drug treatment. Our system applies NLP techniques that capture relationships between anomalies and drug responses from free text. In the field of precision medicine, such relationship will be valuable for determining the right personalized treatment based on patient’s molecular profile. We evaluated our system on different datasets to test the extraction capabilities of the system. It achieved high precision and recall (0.95 and 0.86, respectively) on finding the intended relationships from text. As it is also important to minimize valuable curation time by rejecting non-relevant articles, we evaluated our system’s ability to distinguish relevant articles from non-relevant ones. eGARD achieved true negative rate (TNR) of 0.97, without compromising recall (0.74) as well. Using an external dataset (from PharmGKB project), we showed that eGARD is also able to automatically reproduce 77% of manually curated existing data. After the automatic extraction, eGARD normalizes the gene and disease instances to Entrez and Disease Ontology (DO) IDs, respectively, that will allow for more effective searching as well as enable the use of DO’s hierarchical structure. eGARD extracts additional information that augments the relation extraction and helps a biocurator or a researcher to prioritize curation and determine the confidence or evidence level of extraction.

The robustness and scalability of eGARD were illustrated by applying it to extract relations between genomic anomalies and drug responses from a large set of MEDLINE abstracts. The results are generated in JSON (JavaScript Object Notation) format. JSON is a popular and widely-accepted data-interchange format, that is easy enough for humans to interpret and very convenient for machines to parse. That is why we chose JSON format which will allow for easy exchange of information and integration with other text mining tools and curation frameworks. The next steps for this work would be to apply eGARD on entire MEDLINE, develop a database to house the extracted results along with pointers to the sentences from which they were extracted, optimize a ranking system to classify relevant abstracts that uses the relationships and other entities extracted by eGARD and develop an interactive web interface for curators and researchers to search, manipulate and analyze results. Such developments are beyond the scope of the text mining work described in this paper. Finally, we believe that the extracted information relating genomic anomalies to drug responses will enable researchers to readily generate hypotheses for new precision medicine based clinical trials.

## Acknowledgement

We would like to acknowledge Dr. Simina Boca, Dr. Robert Beckman, Dr. Vishakha Sharma and Dr. Karen Ross for providing valuable use cases and feedback that were helpful in developing and improving eGARD. We gladly acknowledge Samir Gupta for helping with typing of phrases.

## Supporting information

S1 File. List of all patterns used to identify associations between genomic anomalies and drug responses and list of patterns to detect drugs used in treatments.

S2 File. List of these seven gene-drug combinations.

S1 Dataset. InHouseSet1 dataset used for evaluation of eGARD.

S2 Dataset. InHouseSet2 dataset used for evaluation of eGARD.

S3 File. List of FDA-approved drugs for 13 genes in cancer study.

S3 Dataset. List of PMIDs of the PharmGKB dataset.

## References

1. Karapetis CS, Khambata-Ford S, Jonker DJ, O’Callaghan CJ, Tu D, Tebbutt NC, et al. K-ras mutations and benefit from cetuximab in advanced colorectal cancer. N Engl J Med. 2008;359(17):1757–65.

2. Amado RG, Wolf M, Peeters M, Van Cutsem E, Siena S, Freeman DJ, et al. Wild-type KRAS is required for panitumumab efficacy in patients with metastatic colorectal cancer. J Clin Oncol. 2008;26(10):1626–34.

3. Uemura M, Kim HM, Hata T, Sakata K, Okuyama M, Takemoto H, et al. First-line cetuximab-based chemotherapies for patients with advanced or metastatic KRAS wild-type colorectal cancer. Mol Clin Oncol. 2016;5(2):375–9.

4. Marty M, Cognetti F, Maraninchi D, Snyder R, Mauriac L, Tubiana-Hulin M, et al. Randomized phase II trial of the efficacy and safety of trastuzumab combined with docetaxel in patients with human epidermal growth factor receptor 2-positive metastatic breast cancer administered as first-line treatment: the M77001 study group. J Clin Oncol. 2005;23(19):4265–74.

5. Cameron D, Piccart-Gebhart MJ, Gelber RD, Procter M, Goldhirsch A, de Azambuja E, et al. 11 years’ follow-up of trastuzumab after adjuvant chemotherapy in HER2-positive early breast cancer: final analysis of the HERceptin Adjuvant (HERA) trial. Lancet. 2017;389(10075):1195–205.

6. Romond EH, Perez EA, Bryant J, Suman VJ, Geyer CE, Jr., Davidson NE, et al. Trastuzumab plus adjuvant chemotherapy for operable HER2-positive breast cancer. N Engl J Med. 2005;353(16):1673–84.

7. Larkin J, Ascierto PA, Dreno B, Atkinson V, Liszkay G, Maio M, et al. Combined vemurafenib and cobimetinib in BRAF-mutated melanoma. N Engl J Med. 2014;371(20):1867–76.

8. McArthur GA, Chapman PB, Robert C, Larkin J, Haanen JB, Dummer R, et al. Safety and efficacy of vemurafenib in BRAF(V600E) and BRAF(V600K) mutation-positive melanoma (BRIM-3): extended follow-up of a phase 3, randomised, open-label study. Lancet Oncol. 2014;15(3):323–32.

9. Ding PN, Lord SJ, Gebski V, Links M, Bray V, Gralla RJ, et al. Risk of Treatment-Related Toxicities from EGFR Tyrosine Kinase Inhibitors: A Meta-analysis of Clinical Trials of Gefitinib, Erlotinib, and Afatinib in Advanced EGFR-Mutated Non-Small Cell Lung Cancer. J Thorac Oncol. 2017;12(4):633–43.

10. Mok TS, Wu YL, Thongprasert S, Yang CH, Chu DT, Saijo N, et al. Gefitinib or carboplatin-paclitaxel in pulmonary adenocarcinoma. N Engl J Med. 2009;361(10):947–57.

11. Sullivan I, Planchard D. Next-Generation EGFR Tyrosine Kinase Inhibitors for Treating EGFR-Mutant Lung Cancer beyond First Line. Front Med (Lausanne). 2016;3:76.

12. Rao S, Beckman RA, Riazi S, Yabar CS, Boca SM, Marshall JL, et al. Quantification and expert evaluation of evidence for chemopredictive biomarkers to personalize cancer treatment. Oncotarget. 2016.

13. Li MM, Datto M, Duncavage EJ, Kulkarni S, Lindeman NI, Roy S, et al. Standards and Guidelines for the Interpretation and Reporting of Sequence Variants in Cancer: A Joint Consensus Recommendation of the Association for Molecular Pathology, American Society of Clinicalncology, and College of American Pathologists. J Mol Diagn. 2017;19: 4–23.

14. Carr TH, McEwen R, Dougherty B, Johnson JH, Dry JR, Lai Z, et al. Defining actionable mutations for oncology therapeutic development. Nat Rev Cancer. 2016;16(5):319–29.

15. Rehm HL, Berg JS, Brooks LD, Bustamante CD, Evans JP, Landrum MJ, et al. ClinGen-the Clinical Genome Resource. N Engl J Med. 2015;372(23):2235–42.

16. Landrum MJ, Lee JM, Riley GR, Jang W, Rubinstein WS, Church DM, et al. ClinVar: public archive of relationships among sequence variation and human phenotype. Nucleic Acids Res. 2014;42(Database issue):D980–5.

17. Levy MA, Lovly CM, Horn L, Naser R, Pao W. My Cancer Genome: Web-based clinical decision support for genome-directed lung cancer treatment. Journal of Clinical Oncology. 2011;29(15).

18. Griffith M, Spies NC, Krysiak K, McMichael JF, Coffman AC, Danos AM, et al. CIViC is a community knowledgebase for expert crowdsourcing the clinical interpretation of variants in cancer. Nat Genet. 2017;49(2):170–4.

19. UniProt C. Activities at the Universal Protein Resource (UniProt). Nucleic Acids Res. 2014;42(Database issue):D191–8.

20. Wu TJ, Shamsaddini A, Pan Y, Smith K, Crichton DJ, Simonyan V, et al. A framework for organizing cancer-related variations from existing databases, publications and NGS data using a High-performance Integrated Virtual Environment (HIVE). Database (Oxford). 2014;2014:bau022.

21. Amberger J, Bocchini CA, Scott AF, Hamosh A. McKusick’s Online Mendelian Inheritance in Man (OMIM). Nucleic Acids Res. 2009;37(Database issue):D793–6.

22. Beroud C, Hamroun D, Collod-Beroud G, Boileau C, Soussi T, Claustres M. UMD (Universal Mutation Database): 2005 update. Hum Mutat. 2005;26(3):184–91.

23. Thorisson GA, Lancaster O, Free RC, Hastings RK, Sarmah P, Dash D, et al. HGVbaseG2P: a central genetic association database. Nucleic Acids Res. 2009;37(Database> issue):D797–802.

24. Singh A, Olowoyeye A, Baenziger PH, Dantzer J, Kann MG, Radivojac P, et al. MutDB: update on development of tools for the biochemical analysis of genetic variation. Nucleic Acids Res. 2008;36(Database> issue):D815–9.

25. Sherry ST, Ward MH, Kholodov M, Baker J, Phan L, Smigielski EM, et al. dbSNP: the NCBI database of genetic variation. Nucleic Acids Res. 2001;29(1):308–11.

26. Whirl-Carrillo M, McDonagh EM, Hebert JM, Gong L, Sangkuhl K, Thorn CF, et al. Pharmacogenomics knowledge for personalized medicine. Clin Pharmacol Ther. 2012;92(4):414–7.

27. Fleuren WWM, Alkema W. Application of text mining in the biomedical domain. Methods. 2015;74: 97–106.

28. Zhu F, Patumcharoenpol P, Zhang C, Yang Y, Chan J, Meechai A, et al. Biomedical text mining and its applications in cancer research. J Biomed Inform. 2013;46: 200–211.

29. Wei CH, Kao HY, Lu Z. PubTator: a web-based text mining tool for assisting biocuration. Nucleic Acids Res. 2013;41(Web Server issue):W518–22.

30. Wei CH, Kao HY, Lu Z. GNormPlus: An Integrative Approach for Tagging Genes, Gene Families, and Protein Domains. Biomed Res Int. 2015;2015:918710.

31. Leaman R, Islamaj Dogan R, Lu Z. DNorm: disease name normalization with pairwise learning to rank. Bioinformatics. 2013;29(22):2909–17.

32. Leaman R, Wei CH, Lu Z. tmChem: a high performance approach for chemical named entity recognition and normalization. J Cheminform. 2015;7(Suppl 1 Text mining for chemistry and the CHEMDNER track):S3.

33. Wei CH, Harris BR, Kao HY, Lu Z. tmVar: a text mining approach for extracting sequence variants in biomedical literature. Bioinformatics. 2013;29(11):1433–9.

34. Wei CH, Kao HY, Lu Z. SR4GN: a species recognition software tool for gene normalization. PLoS One. 2012;7(6):e38460.

35. Singhal A, Simmons M, Lu Z. Text mining for precision medicine: automating disease-mutation relationship extraction from biomedical literature. J Am Med Inform Assoc. 2016;23(4):766–72.

36. Singhal A, Simmons M, Lu Z. Text Mining Genotype-Phenotype Relationships from Biomedical Literature for Database Curation and Precision Medicine. PLoS Comput Biol. 2016;12(11):e1005017.

37. Hakenberg J, Voronov D, Nguyen VH, Liang S, Anwar S, Lumpkin B, et al. A SNPshot of PubMed to associate genetic variants with drugs, diseases, and adverse reactions. J Biomed Inform. 2012;45(5):842–50.

38. Xu R, Wang Q. A knowledge-driven conditional approach to extract pharmacogenomics specific drug-gene relationships from free text. J Biomed Inform. 2012;45(5):827–34.

39. Rance B, Doughty E, Demner-Fushman D, Kann MG, Bodenreider O. A mutation-centric approach to identifying pharmacogenomic relations in text. J Biomed Inform. 2012;45(5):835–41.

40. Rinaldi F, Schneider G, Clematide S. Relation mining experiments in the pharmacogenomics domain. J Biomed Inform. 2012;45(5):851–61.

41. Pakhomov S, McInnes BT, Lamba J, Liu Y, Melton GB, Ghodke Y, et al. Using PharmGKB to train text mining approaches for identifying potential gene targets for pharmacogenomic studies. J Biomed Inform. 2012;45(5):862–9.

42. Garten Y, Altman RB. Pharmspresso: a text mining tool for extraction of pharmacogenomic concepts and relationships from full text. BMC Bioinformatics. 2009;10 Suppl 2:S6.

43. Garten Y, Coulet A, Altman RB. Recent progress in automatically extracting information from the pharmacogenomic literature. Pharmacogenomics. 2010;11(10):1467–89.

44. Coulet A, Cohen KB, Altman RB. The state of the art in text mining and natural language processing for pharmacogenomics. Journal of Biomedical Informatics. 2012;45(5):825–6.

45. Mahmood AS, Wu TJ, Mazumder R, Vijay-Shanker K. DiMeX: A Text Mining System for Mutation-Disease Association Extraction. PLoS One. 2016;11(4):e0152725.

46. Gupta S, Ross KE, Tudor CO, Wu CH, Schmidt CJ, Vijay-Shanker K. miRiaD: A Text Mining Tool for Detecting Associations of microRNAs with Diseases. J Biomed Semantics. 2016;7(1):9.

47. Schwartz AS, Hearst MA. A simple algorithm for identifying abbreviation definitions in biomedical text. Pac Symp Biocomput. 2003:451–62.

48. Peng Y, Tudor,C., Torii,M., Wu,C.H., Vijay-Shanker,K., editor iSimp: A Sentence Simplification System for Biomedical Text. In Proceedings of the 2012 IEEE International Conference on Bioinformatics and Biomedicine; 2012.

49. Narayanaswamy M, Ravikumar KE, Vijay-Shanker K. A biological named entity recognizer. Pac Symp Biocomput. 2003:427–38.

50. Kibbe WA, Arze C, Felix V, Mitraka E, Bolton E, Fu G, et al. Disease Ontology 2015 update: an expanded and updated database of human diseases for linking biomedical knowledge through disease data. Nucleic Acids Res. 2015;43(Database> issue):D1071–8.

51. Ding R, Arighi CN, Lee JY, Wu CH, Vijay-Shanker K. pGenN, a gene normalization tool for plant genes and proteins in scientific literature. PLoS One. 2015;10(8):e0135305.

